# Comprehensive functional screening of taste sensation *in vivo*

**DOI:** 10.1101/371682

**Authors:** Jisoo Han, Myunghwan Choi

## Abstract

The initial event in taste sensation is mediated by taste cells on the tongue that translate ingested chemicals into cellular signals. Current understanding on this cellular level taste encoding process has relied on *ex vivo* model systems that cannot fully recapitulate natural cellular microenvironment *in vivo*. To resolve this methodological limitation, we invented a microfluidics-on-a-tongue imaging chamber that has integrated multichannel microfluidics for auto-controlled tastant delivery. Using this system, we screened over 100 fungiform taste cells to the five basic taste qualities, and obtained comprehensive functional maps. We revealed that taste cells are composed of 70% of single-tuned and 30% of dual-tuned cells, and also discovered a novel population of dual-tuned taste cells encoding a positive valence. We believe that our novel screening platform will pave a way for the deeper understanding of taste coding logic.

## INTRODUCTION

Taste is fundamental to life, as it guides the intake of nutritious contents and the avoidance of toxic chemicals in foods (Lindemann, 2001). In mammals, the foremost sensor is taste cells on the tongue that convert ingested chemicals into cellular activity (Chaudhari and Roper, 2010). The cellular information is subsequently conveyed to the central nervous system (Finger, 2005). The understanding of this encoding-decoding process is one of the fundamental questions in the field of taste (Liman et al., 2014; Roper and Chaudhari, 2017).

There has been decades of studies on how taste information is encoded into taste cells using various tools such as electrophysiology (Yoshida et al., 2010), genetics (Chandrashekar et al., 2006; Lee et al., 2017; Ohmoto et al., 2013), and microscopy (Chandrashekar et al., 2010; Dvoryanchikov et al., 2011; Oka et al., 2013; Tomchik et al., 2007). However, all of these existing tools inevitably involve invasive sample preparations for measuring functional activities of taste cells: digestion of extracellular matrices (Chandrashekar et al., 2010), isolation of individual taste cells or buds (Yoshida et al., 2006), and incubation in artificial environments. Evidently, these *ex vivo* model systems cannot fully recreate the native cellular microenvironment. For example, the physiological protective barriers formed at intercellular junctions are degraded (Holland, V.F., Zampighi, G.A., Simon, 1991), blood perfusion mediating endocrine signaling is lost (Wu et al., 2015), and feedback signals from afferent nerve fibers are lost (Vandenbeuch and Kinnamon, 2016). Consequently, the *ex vivo* condition severely interferes with the reliable functional screening of multiple tastants requiring hours of observation. The first imaging window for observing taste cells in a living mouse has previously been reported, but it lacked critical functionalities such as stable real-time imaging and controlled tastant delivery (Choi et al., 2015).

Here, we report a novel image-based screening platform, named “microfluidics-on-a-tongue” that enables high-throughput functional screening of taste cells *in vivo*. By introducing the microfluidics-based tastant delivery and ratiometric calcium imaging, this system provides unprecedented reliable real-time activity of taste cells in their native environment. We exploited this platform to demonstrate a comprehensive *in vivo* screening of the five basic tastes (sweet, umami, salty, sour and bitter) in over 100 taste cells. Through this screening, we reveal novel populations of taste cells that are tuned to dual taste qualities.

## RESULTS

### System design and validation

The microfluidics-on-a-tongue system was designed to provide stable microscopic imaging of the dorsal surface of the anterior tongue (i.e., fungiform taste bud) while controllably delivering multiple liquid-phase tastants (Fig. 1). The main device was custom-made with stainless steel to provide mechanical rigidity in a small form factor, as well as provide corrosion resistance for long-term usage. The device is composed of two units, the top and bottom units that sandwich the mouse tongue to stabilize the physiological motions caused by the heartbeat and breathing (Fig. 1a,b; Supplementary Fig. 1). The bottom unit holds the externalized tongue by chemically gluing the ventral part while the top unit is placed at an adjustable height of ~1.5 mm from the bottom unit to gently stabilize the tongue (Supplementary Fig. 2). To enable cellular-level imaging, the top unit contains an optical window made with a circular cover slip. An opening of 3 mm in diameter covers nearly half of the anterior tongue. This window size is compatible with an uninterrupted high-NA volumetric imaging of the taste buds lying ~0.8 mm from the surface of the device (Fig. 1c) (Beaurepaire and Mertz, 2002). In addition, the top unit integrates a spindle-shaped microfluidic channel for the controlled delivery of multiple liquid-phase tastants. The input port is connected to an 8-channel micro-manifold that is fed by a computer-controlled perfusion system. In our initial trials, we observed that the fluid flow generates severe tissue motions, especially while switching the fluidic input channels. To stabilize the flow, we connected the output port to a syringe pump, clamping the fluid drainage at a fixed flow rate of 300 μl∙min^-1^. By balancing the input and output flow rates, we could obtain a stable quasi-steady-state flow. In addition, we introduced a transparent porous membrane (pore size = 30 μm) at the interface of the tongue and the microfluidic channel to minimize the effect of fluid shear stress. In the control trials with four channels filled with artificial saliva, we obtained microscopic stability even while switching abruptly between input channels (Supplementary Fig. 3).

**Figure 1.**
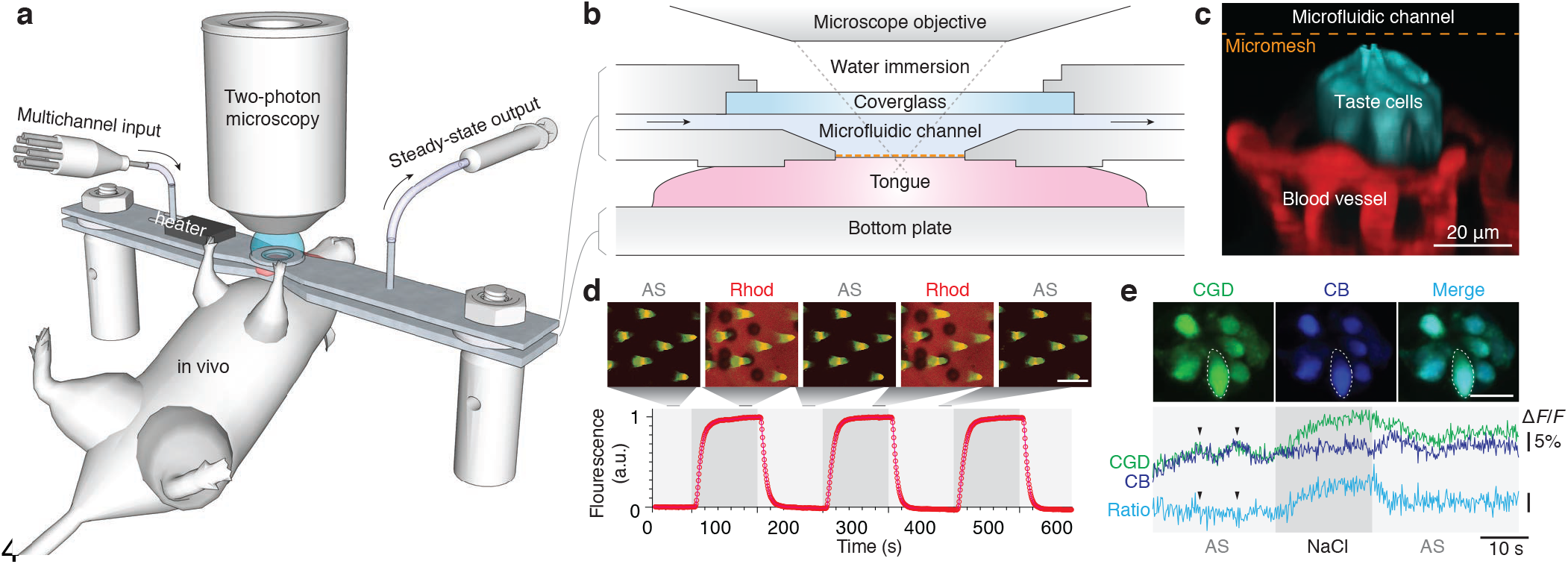
Microfluidics-on-a-tongue platform. (**a**) A schematic diagram. The tongue of the anesthetized mouse was mounted on the microfluidics-on-a-tongue device. Various liquid-phase tastants were delivered through a multichannel input controlled by a computerized fluidic system, while the taste bud was imaged using a two-photon microscope. (**b**) A cross-sectional view of the imaging chamber. Arrows indicate the direction of fluidic stimuli. The orange dashed line indicates the micromesh used to minimize the movement of artifact during fluidic switching. (**c**) A 3-dimensional fluorescent micrograph of the taste bud. The taste cells in cyan were co-labeled with calcium-green dextran (CGD) and cascade blue (CB). The blood vessel in red was labeled with an intravenous rhodamine-dextran. (**d**) Characterization of the microfluidic system. The dorsal surface of the tongue was serially imaged with alternating inputs of artificial saliva (AS) and fluorescent rhodamine (Rhod). The cone-shaped filiform papillae (green) were made visible by keratin autofluorescence. The fluorescent traces were fitted to exponential curves. Scalebar, 100 μm. (**e**) Ratiometric calcium imaging. Taste cells loaded with both a calcium-sensitive dye (CGD) and a calcium-insensitive dye (CB) were imaged at ~ 6 Hz with salty taste stimulus (400 mM NaCl). Note that ratiometric analysis effectively eliminates the artefactual fluctuations (arrowheads) while preserving the taste-evoked calcium response. Scalebar, 20 μm.

After establishing the stabilized condition, we characterized the fluidic parameters that determine the kinetics of taste stimuli to the taste cells. To achieve this, we alternated between fluidic input channels filled with either control or fluorescent artificial saliva, while performing real-time microscopy on the apical surface of the taste bud (Fig. 1d). The infusion and washout kinetics followed one-phase exponential curves with a time constant of 8.53±1.6 s. The latency between switching on a channel and the arrival of the tastant to the taste bud was 6.49±0.29 s. Based on this information, we designed the parameters for tastant delivery and analyzed the functional data, as described in Online Methods.

When tested with various physiologic tastants, however, we noted a critical issue caused by the difference in the refractive indices of the solutions (Supplementary Table 1). Artificial saliva has a refractive index close to water (1.333) but the additional solutes in the taste solutions proportionally increase the refractive index (Supplementary Fig. 4). For example, an umami solution prepared with 100 mM MPG has a higher refractive index of 1.336 (Δn = 0.003) that introduces considerable optical aberrations, and thus artefactual signals. This optical noise often overwhelms fluorescent calcium imaging that typically depends on a small change in fluorescence intensity (Δ*F*/*F* ≈ 1–10%). To resolve this issue, we adopted a ratiometric calcium imaging that quantifies the relative change in calcium-sensitive dye to calcium-insensitive dye (Fig. 1e) (Nguyen et al., 2016; Thestrup et al., 2014). We chose calcium-green dextran (CGD) for the calcium-sensitive dye and cascade blue (CB) for the calcium-insensitive dye due to their anionic charge, two-photon efficiency, and minimal spectral overlap. To our surprise, the ratiometric approach cleanly resolved the refractive index issue because the optical aberration affects the fluorescence intensities of both dyes to the same degree. Furthermore, it effectively suppressed system-level optical noises such as laser power drift that is typically ±0.5% in optical intensity, leading to ±1% noise increase in two-photon fluorescence (Supplementary Fig. 4).

Finally, we investigated if physiological blood perfusion to the taste bud is maintained in our preparation (Supplementary Fig. 5). To measure the blood flow, we intravenously administered a fluorescent dextran and traced negatively-stained blood cells in the capillaries surrounding the taste bud (Shih et al., 2012). By measuring the blood flow while adjusting the height of the top unit, we revealed that compressive strain of over 35% on the tongue near-completely ceases the blood flow. As the ischemic condition often leads to irreversible calcium response or inactive state of taste cells (Szydlowska and Tymianski, 2010), we set the compressive strain to be less than 10% and confirmed that blood flow was intact before every imaging session.

### Comprehensive functional screening

Having validated the system, we examined how taste cells are tuned to different taste qualities. We exposed a series of tastants at physiological concentrations to the tongue with intervals of 4 min while performing serial two-photon imaging of the taste bud at ~6 Hz. For each taste cell, the intensity trace for calcium-sensitive dye was divided by that for calcium-insensitive dye, and the resulting ratiometric trace denoted as the relative change to the baseline (Δ*F*/*F*). In this protocol, the responsive cells showed robust activation through repeated stimulations (Fig. 2a,b). For each taste bud, we typically screened 4-8 stained taste cells (lower and upper quartiles) to the five basic tastes, with 4 repeated stimuli for each taste quality (Fig. 2c). Cells were classified as responsive to the taste quality if statistically significant positive responses were observed in sync with the tastant delivery. We presented the spatial profile of the cellular responses into a taste map for each taste bud (Fig. 2d). In total, we screened 101 cells in 17 taste buds (n = 17 mice).

**Figure 2.**
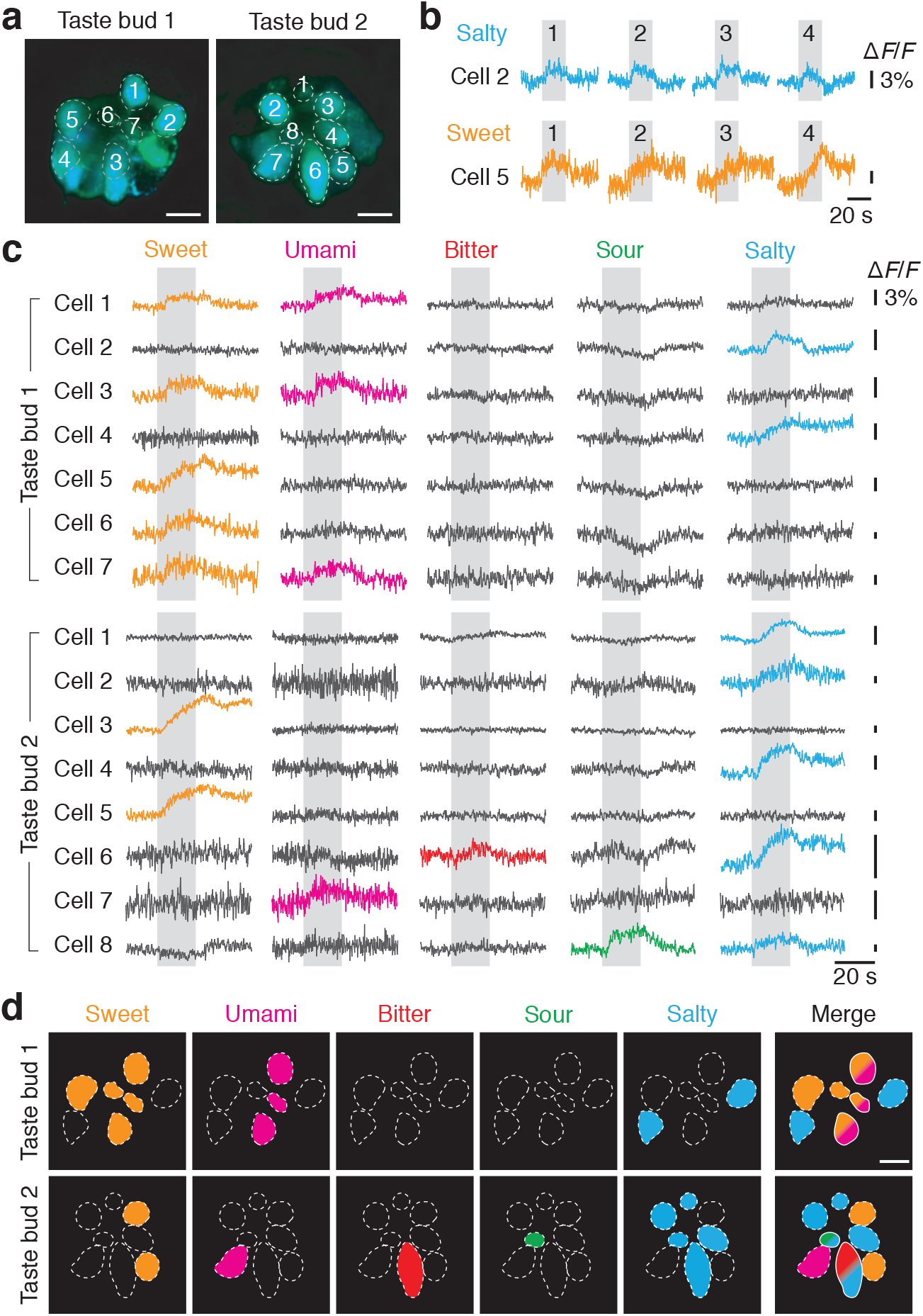
Comprehensive functional mapping of the fungiform taste buds *in vivo*. (**a**) Fluorescence images of the taste buds loaded with the ratiometric dyes. Dashed lines demarcate each numbered taste cell. Scalebar, 30 μm. (**b**) Repeatability of taste-evoked calcium response. In taste bud 1, cells 2 and 5 repeatedly responded four times to salty (400 mM NaCl) and sweet (40 mM acesulfame potassium) stimuli, respectively. The shaded regions represent the duration of the taste stimuli. (**c**) Calcium response of each taste cell to the five basic taste stimuli. Each trace is the average of repeated trials. Colored traces are classified as “responsive” (p < 0.01). (**d**) Taste map. For each taste quality, responsive cells from the calcium traces in (c) were colored. In the merged image, single-colored and dual-colored cells represent single-tuned and dual-tuned cells, respectively. Scalebar, 10 μm.

In the screening data, we observed that ~70% of fungiform taste cells were tuned to a single taste quality, supporting the labeled line model that each taste quality is encoded by separate taste cells (Fig. 3a) (Yarmolinsky et al., 2009). In single-tuned cells, the cell population for each taste quality was unevenly distributed: 30% salty, 28% bitter, 24% sweet, 11% sour, and 7% umami. Interestingly, we observed two “bitter taste buds” (out of 17 taste buds) composed of over 85% of bitter-responding cells (Supplementary Fig. 6). Other than the two bitter taste buds, we did not observe notable topographical clustering of functional cell types within a taste bud.

**Figure 3.**
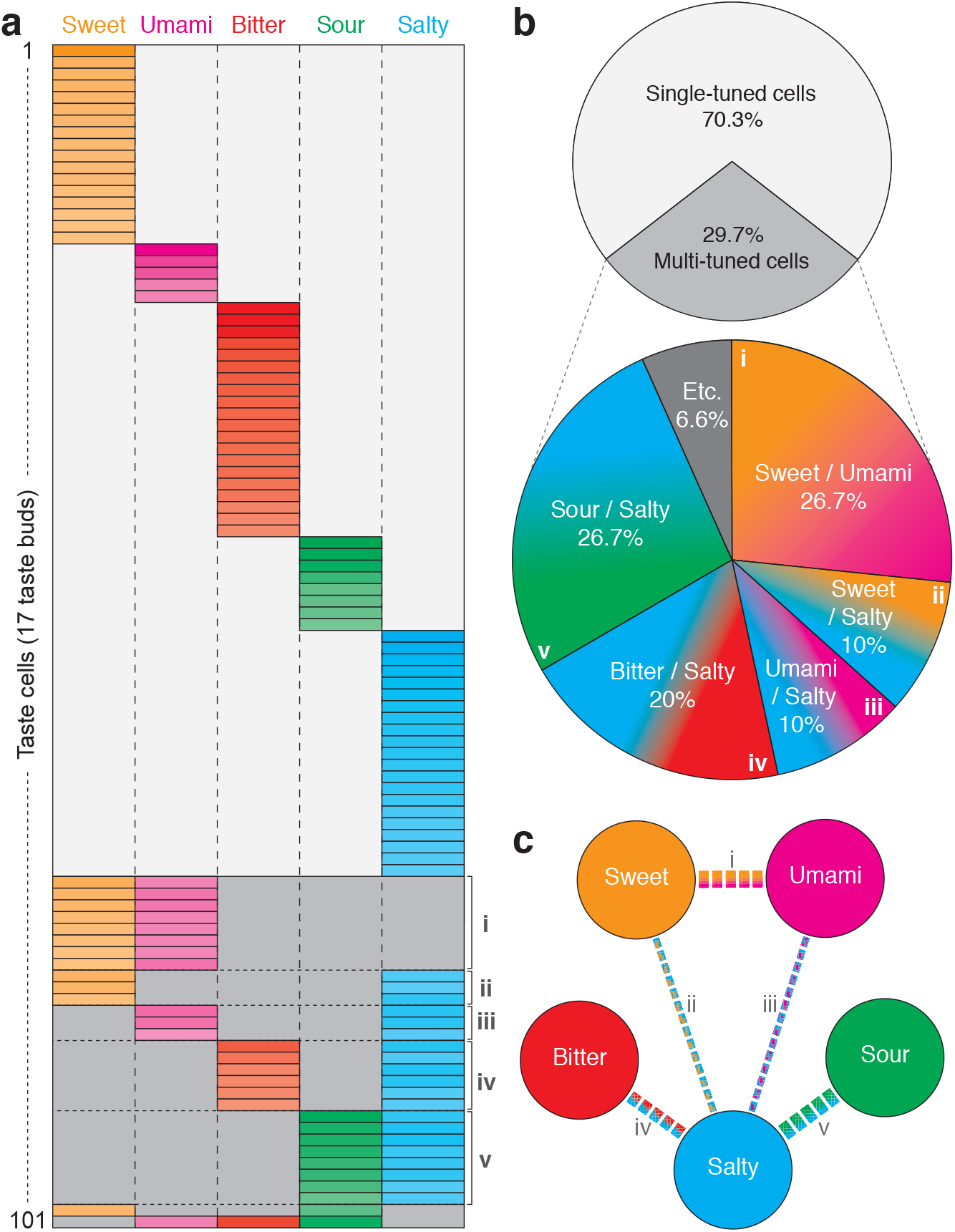
Tuning of fungiform taste cells. (**a**) Summary of taste-evoked cellular responses. Each row represents a taste cell (101 cells from 17 taste buds) while each colored region indicates the statistically significant calcium responses from 4 repeated trials (p < 0.01). The color intensity represents the magnitude of the calcium response (Δ*F*/*F*). (**b**) Summary statistics of taste-evoked cellular responses. Note that 30% of taste cells were multi-tuned cells, with most of them being dual-tuned. (**c**) Graphical representation of connectivity among the five basic tastes. Thicknesses of the edges indicate strengths of the connectivity. Note that sweet and umami show a strong connection while salty is connected with all the other taste qualities.

Yet, ~30% of taste cells were tuned to multiple taste qualities, mostly to dual taste qualities (Fig. 3b). Among 10 possible combinations (_5_C_2_) of dual-tuned responses (Chandrashekar et al., 2010), we discovered that only several well-defined modes of inter-taste crosstalk exists (Fig. 3c). Notably, there was strong coupling between sweet and umami, the two appetitive taste qualities. Although recent studies have reported the sweet-umami crosstalk at the level of downstream afferent neurons (Barretto et al., 2015; Wu et al., 2015), our finding provides the first direct evidence of the functional crosstalk present in peripheral taste cells. We also discovered that salty tastes exhibit functional connections to all other basic tastes, either preferred (sweet and umami) or aversive (bitter and sour) tastes. This result may be attributed to the unique bimodal behavior of salty taste – preferred at low concentrations (<150 mM) but avoided at high concentrations (>300 mM) (Chandrashekar et al., 2010).

### Verification of sweet-umami cells

We further investigated the molecular mechanism of the sweet-umami cells responding to the two different taste qualities. Sweet and umami taste receptors are formed by hetero-dimerization of the three T1R family: T1R2/T1R3 for sweet and T1R1/T1R3 for umami (Zhao et al., 2003). Therefore, we hypothesized that these cells may express all the three T1R gene family to be responsive to both sweet and umami taste qualities. Previous studies on this issue resulted in controversial outcomes presumably because only two mRNAs were detected for each tissue section (Kim et al., 2003; Nelson et al., 2001). To clear this controversy, we performed 3-color fluorescent *in situ* hybridization (T1R1, T1R2 and T1R3) on lingual slices with single-molecular sensitivity and counted the single-mRNAs for each fungiform taste cell (Fig. 4) (Wang et al., 2012). To reliably demarcate individual taste cells, we counter-stained the nuclei with DAPI and took a differential-interference contrast (DIC) image (Fig. 4a,b). Consistent with our functional screening data, we discovered that ~10% of taste cells transcribe all the three T1R gene family that are potentially sweet-umami cells (Fig. 4c,d).

**Figure 4.**
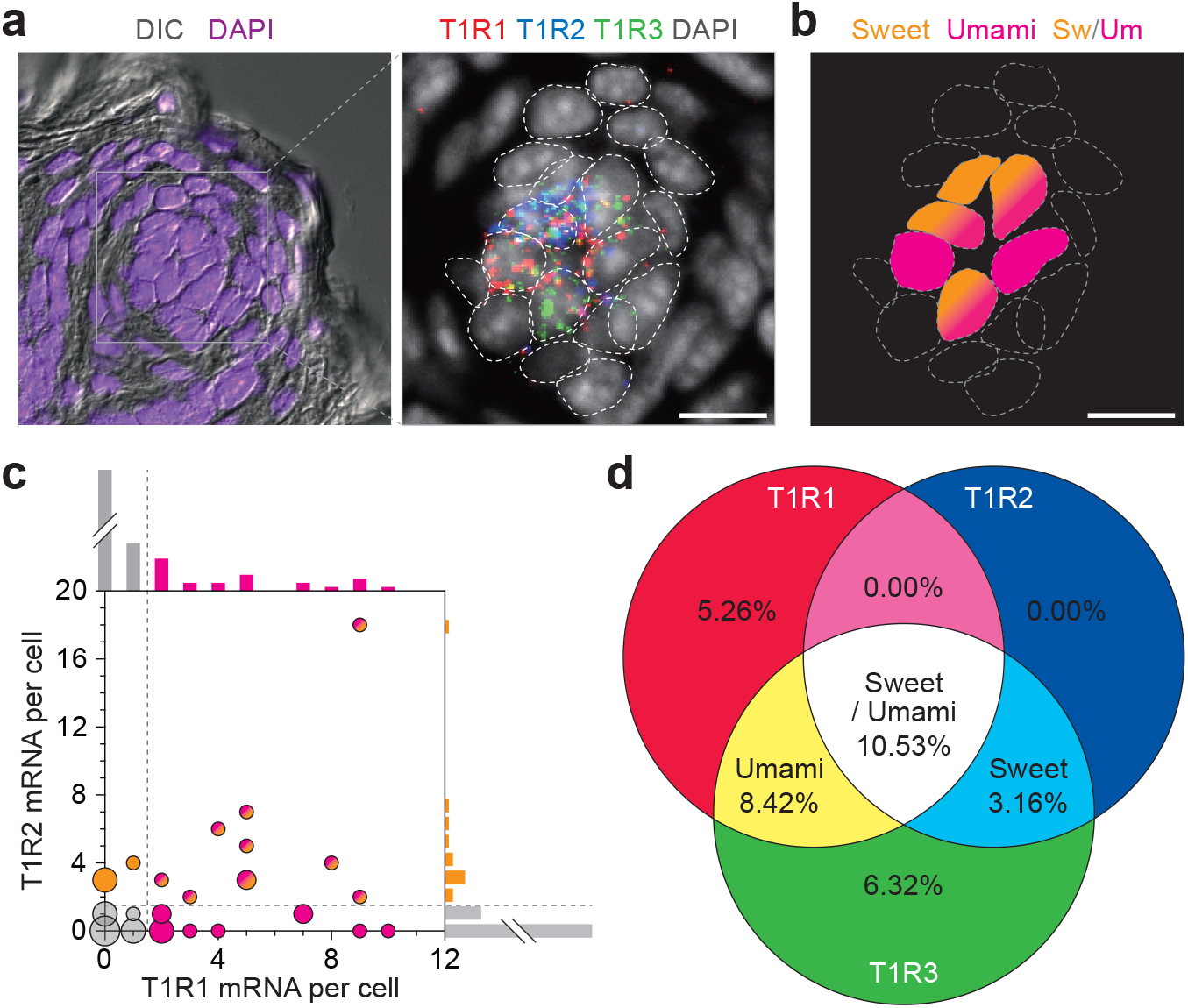
Transcriptional verification of sweet-umami cells. (**a**) Triple-color fluorescent *in situ* hybridization of T1R family (T1R1, T1R2 and T1R3). Differential interference contrast (DIC) and nuclear stain (DAPI) were imaged to delineate the margins for cells and the nucleus. On the right panel, each fluorescent dot represents a single mRNA of T1R1 (red), T1R2 (blue), or T1R3 (green), with each nuclear margin demarcated with dashed lines. Scalebar, 10 μm. (**b**) Map of sweet (T1R2/T1R3) and umami (T1R1/T1R3) cells based on mRNA profile. Scalebar, 10 μm. (**c**) Single-cell mRNA expression for T1R1 and T1R2 (n = 95 cells). The relative size of the circle represents the number of cells. Dashed lines indicate the threshold level, corresponding to “>1 mRNA per cell”. (**d**) Venn diagram of T1Rs transcriptome. Note that ~10% of taste cells transcribe to both sweet and umami receptors, consistent with our functional screening shown in Fig. 3.

## DISCUSSION

We have reported a novel microfluidics-on-a-tongue platform that offers reliable functional imaging of fungiform taste cells in their native living state, for the first time. Harnessing this platform, we comprehensively screened ~100 taste cells to the five basic taste qualities, and obtained functional classification of the taste cells, composed of 70% of single-tuned and 30% of dual-tuned cells. We further discovered that the fungiform taste bud contains ~10% of sweet-umami cells, potentially encoding a positive valence.

Our *in vivo* screening result may resolve long-standing controversies on the breadth of tuning at the taste bud. Previous studies performed *ex vivo* resulted in controversial outcomes. Several studies reported predominant multi-tuned cells responding to up to five different taste qualities (Caicedo et al., 2002; Tomchik et al., 2007), while other studies reported mostly single-tuned cells with minimal crosstalk (Chandrashekar et al., 2006). Our comprehensive screening clarifies that there exist ~70% of single-tuned cells and ~30% of dual-tuned cells and the dual-tuned cells exhibit well-defined modes of crosstalk, composed of sweet-umami and salty-others. The previously reported crosstalk between high salt and aversive taste qualities (i.e., bitter and sour) were also found within the modes of the crosstalk we observed, suggesting the validity of our findings (Oka et al., 2013).

Interestingly, our screening result on taste cells bears resemblance with recent *in vivo* studies on geniculate neurons that relay taste information from the fungiform taste bud to the brain (Barretto et al., 2015; Wu et al., 2015). First, we revealed that a novel population of taste cells that respond to both sweet and umami comprises ~10% of the total population. Similarly, geniculate neurons showed that 3–11% of the total population responds to both sweet and umami. Moreover, ~45% of geniculate neurons that responded to umami also responded to sweet. This also agrees well with our result on taste cells (~47%). These sweet-umami neurons were previously conceived as a convergence of taste information from distinct sweet and umami taste cells but our finding provides an alternative possibility that convergence of sweet and umami (conversely, positive valence) occurs at the earliest taste encoding process (Barretto et al., 2015). The gustatory cortex consistently encodes the sweet and umami in closely apposed cortical fields (Chen et al., 2011), and a behavioral assay based on conditioned taste aversion demonstrated considerable cross-generalization between the two appetitive taste qualities (Heyer et al., 2004). Furthermore, our finding that salty taste is connected to all other taste qualities was also observed in the geniculate ganglion. This may be attributed to the unique bimodal behavior of salty taste: preferred at lower concentrations (<150 mM) but avoided at higher concentrations (>300 mM). The moderate saltiness used in our screening (~250 mM) might have evoked both preferred (i.e., sweet and umami) and aversive (i.e., bitter and sour) pathways (Chandrashekar et al., 2010; Oka et al., 2013). By contrast, bitter-sour crosstalk that is the most prevalent in geniculate neurons was nearly unobserved in the taste bud. This result suggests that convergence of bitter and sour may arise at the neuronal level. This is consistent with the finding that bitter-sour neurons receive input from bitter T2R-expressing cells (Barretto et al., 2015). We envision that detailed comparative analysis of the screening data along the taste transduction pathway will offer better understanding of information processing on tastes (Dando and Roper, 2009; Huang et al., 2009; Roper, 2006).

From a technical standpoint, our screening platform can be improved further. Although the ratiometric approach provided effective correction of imaging artifacts, it could not correct the large axial motion in the focal plane (>3 μm) caused by high-index tastants (n > 1.34) or the non-rigid tissue motion caused by sour tastes. We also noted that the pH change induced by sour tastants could affect the absorbance or quantum yield of fluorophores. Although our current protocol showed cellular calcium response greater than the artifact, studies on strong acids may require pH-insensitive fluorophores. In addition, the electrophoretic loading of fluorophores can affect cellular functionalities. We indeed observed that voltage levels higher than 2 Vpp could be cytotoxic. Introducing transgenic animals with a genetic calcium indicator (e.g., GCaMPs) in the taste cells would provide an effective solution to this issue and further enable cell-type specific labeling. Finally, two-photon microscopy based on point scanning only captures a small fraction of information from the specimen restricted to a focal plane within a single taste bud. Advanced volumetric scanning systems can capture more spatiotemporal information including axially propagating intracellular calcium signals (Bouchard et al., 2015; Song et al., 2017). Random access scanning with ultrafast acoustic-optic scanners would enable the imaging of multiple taste buds in a single trial for attaining higher throughputs screening (Katona et al., 2012).

The microfluidics-on-a-tongue platform is broadly applicable to other areas of studies on the tongue. This platform may provide quantitative representation of tastes that have only been communicated with subjective description. Taste quantification will have a huge impact on the food industry by enabling objective information transfer between the industry and their customers. There is also increasing interest on drug-taste interaction, as taste change is one of the most common side effects of drugs (Clark et al., 2012). Screening of functional changes by drugs offers a reliable quantification of drug-taste interaction. Similar screening can be utilized to find taste modulatory compounds to enhance preferred tastes or inhibit avoided ones (Zuker, 2015). Homeostatic maintenance of taste cells and age-related change in taste sensation are also interesting fields that can benefit from our technique (Lee et al., 2017).

## MATERIALS AND METHODS

### Animal

Male mice aged 8–12 weeks C57BL/6 (Orient Bio) were used for this research. All animal experiments were performed in compliance with institutional guidelines and approved by the subcommittee for research animal care at Sungkyunkwan University.

### Mouse preparation

Mixture of 0.12 mg/g zoletil (Virbac) and 0.01 mg/g rompun (Bayer Korea) was intraperitoneally injected for mice anesthesia. 2.5% w/v TRITC-dextran (500kDa, Sigma Aldrich) in phosphate-buffered saline (Life technologies) was injected intravenously for blood flow imaging. To reduce movement artifact, the animal skull was fixed; approximately 1 cm^2^ of the skin of the mouse was removed and periosteum wiped out using cotton swab. A metal rod ~3.9 mm in diameter and ~20 mm in length was attached to the mouse skull with dental glue (DENTKIST, South Korea). This rod was tied up to a surgical board using Epoxy putty (PSI, Polymeric System, INC). Afterwards, the mouse tongue was softly drawn using a plastic tweezer. The ventral part of the tongue and lower lip were glued with instant medical adhesive (Loctite 4161, Henkel Corporation) to the bottom unit of the fluidic device for immobility. The exposed surface of the mouse tongue was preserved under wet conditions with artificial saliva.

### Electroporation

Electrophoretic bulk loading was performed for taste cell staining 12-36 h before the experiment using Calcium Green-1 dextran, CGD (3,000 kDa, Invitrogen) and cascade blue dextran, GB (3,000 kDa, Invitrogen). Both dyes were introduced for ratiometric analysis since CGD is calcium-sensitive and GB is calcium-insensitive. The anesthetized mouse tongue was drawn out of the mouth using a pair of blunt plastic forceps. Optical lens paper (7 by 7 mm^2^) was steeped in 5 % w/v CGD and 2.5 % w/v CB with 7:3 volumetric ratio and placed on the anterior surface of the mouse tongue. The tongue was then sandwiched between electrical tweezers, 5 mm in diameter (Tweezertrode, Harvard Apparatus). Electric pulses (voltage: 2 Vpp, frequency: 1 Hz, pulse width: 100 ms, duration: 50–100 s) from a function generator (KEYSIGHT TECHNOLOGIES, 33220A) was used for electrophoresis (Choi et al., 2015).

### Solutions

Artificial saliva (2 mM NaCl, 5 mM KCl, 3 mM NaHCO_3_, 3 mM KHCO_3_, 0.25 mM CaCl_2_, 0.25 mM MgCl_2_, 0.12 mM K_2_HPO_4_, 0.12 mM KH_2_PO_4_, 1.8 mM HCl, pH7) (Danilova, 2003) was used as control state for taste cells and washing out after tastant stimulation. The following were tastants used in this study: Bitter (mixture of 2.5–5 mM quinine, 2.5–5 mM denatonium and 10–20 μM cycloheximide), Sweet (40 mM AceK), Sour (10 mM Citric Acid), Salty (100 mM low sodium, 250 mM or 400 mM high sodium), Umami (mixture of 100 mM MPG and 1 μm IMP).

### Stabilization of the microfluidics-on-a-tongue device

Liquid delivery was controlled using a pressure controller and its software (Octaflow II, ALA Scientific). Suction syringe was connected to prevent perturbation caused by switching channels. Typically 0.4 psi for 8- channel input and 300 μl∙min^-1^ for discharge output was applied to establish the steady-state condition. A stabilized “microfluidic-on-a-tongue-device” was then placed on the surface of the mice tongue. Both ends of the device were screwed tight to prevent fluidic turbulence or physiological movement. Curved washers were inserted between the upper and bottom units to protect the tongue from damage due overpressure.

### Calcium imaging

After the preparation, the mouse was moved to the stage under a two-photon microscope. A heating pad was used for maintaining body temperature. The anterior part of the tongue was exposed to the chamber opening while artificial saliva was constantly supplied to the taste buds and/or cells. Each session was composed of 20 s stimulation and 4 min washing with artificial saliva. During each session, taste cell calcium imaging was captured for about 60-80 s to avoid photo toxicity. A water immersion objective (16X, NA 0.80, Nikon or 20 X, NA 0.92 Leica) was equipped with a two-photon microscope body (Bruker), with 800–810 nm wavelength used for the two-photon excitation (Coherent, Chameleon Vision II). Fluorescent images were captured at ~6 Hz with emissions at 460±25 nm for CB, 525±25 nm for CGD and 605±35 nm for TRITC-dextran.

### In situ hybridization

Cervical dislocation of the anesthetized mouse was performed and the tongue extracted. The dissected mouse tongue was directly dropped in liquid N2 for instant freezing. The frozen tongue was embedded in an optimal cutting temperature (OCT) compound block and kept overnight at −80 °C in a deep freezer. The following day, the OCT block was moved to a −20 °C freezer for one hour and cryosectioned into 10-μm-thick sections. The in situ hybridization was carried out in compliance with the RNAscope procedure. The accession numbers for the mRNAs used in this study is as follows: T1R1 (NM_031867.2), T1R2 (NM_031873.1) and T1R3 (NM_031872.2).

### Statistical analysis

We used MATLAB software for statistical analyses. Paired or unpaired t-tests (two-sided) were used for group comparisons by assuming normality based on previous literatures. Neither randomization nor blinding was applied. For regression analyses, we presented correlation coefficient (R^2^) along with the sample size. The data was presented as the mean ± standard error. We considered *p*-values less than 0.01 to be statistically significant.

### Code availability

The Matlab codes used for the simulation studies are available from the corresponding author upon reasonable request for purposes of academic research.

### Data availability

All relevant data are available from the corresponding author upon reasonable request.

## Supplementary Data

**Supplementary Table 1.**
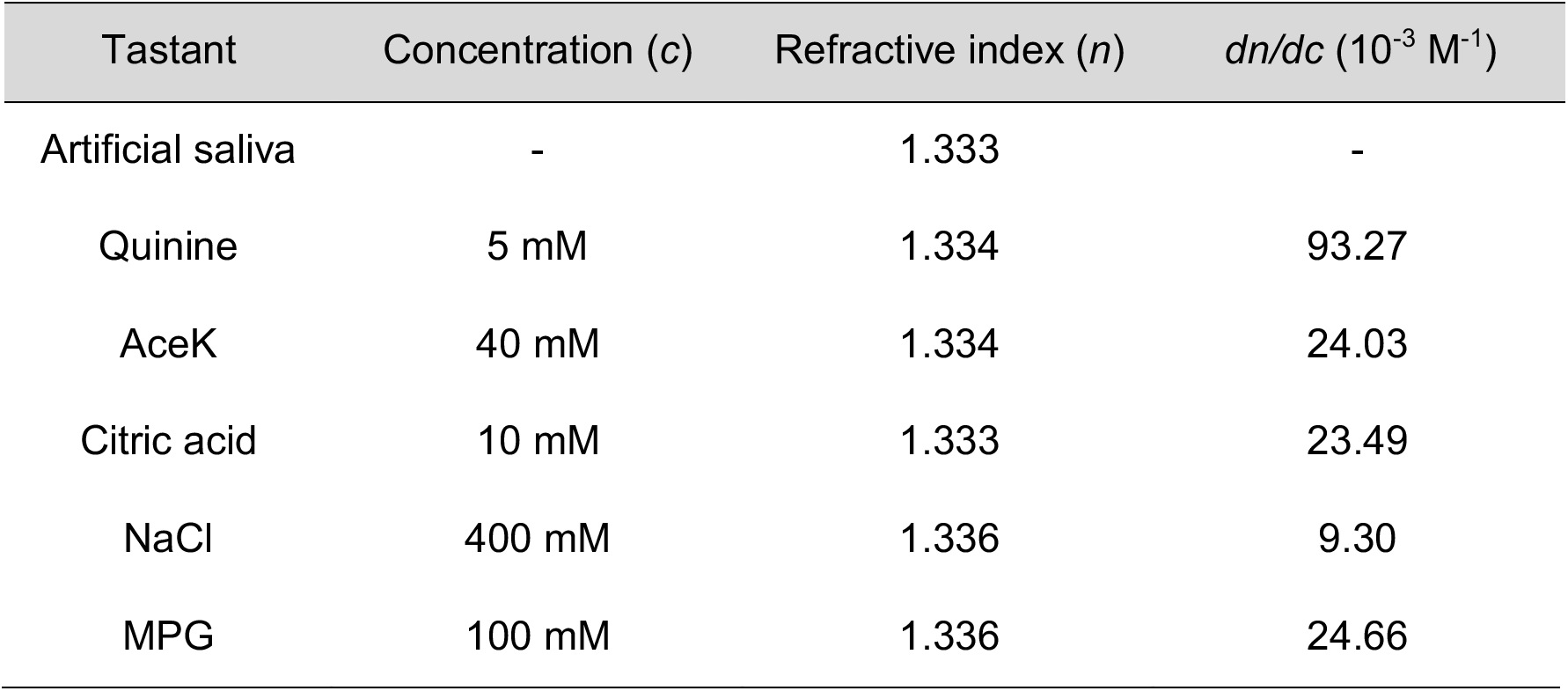
Optical characterization of tastants. Tastants were prepared in artificial saliva and refractive indices were measured at room temperature by a digital refractometer (ORD 85BM, Kern & Sohn). The specific refractive index increment (*dn/dc*) was determined by a linear-fit to data measured at five different concentrations.

| Tastant | Concentration (*c*) | Refractive index (*n*) | $dn/dc$ ($10^{-3} \text{ M}^{-1}$) |
| --- | --- | --- | --- |
| Artificial saliva | - | 1.333 | - |
| Quinine | 5 mM | 1.334 | 93.27 |
| AceK | 40 mM | 1.334 | 24.03 |
| Citric acid | 10 mM | 1.333 | 23.49 |
| NaCl | 400 mM | 1.336 | 9.30 |
| MPG | 100 mM | 1.336 | 24.66 |

**Supplementary Fig. 1.**
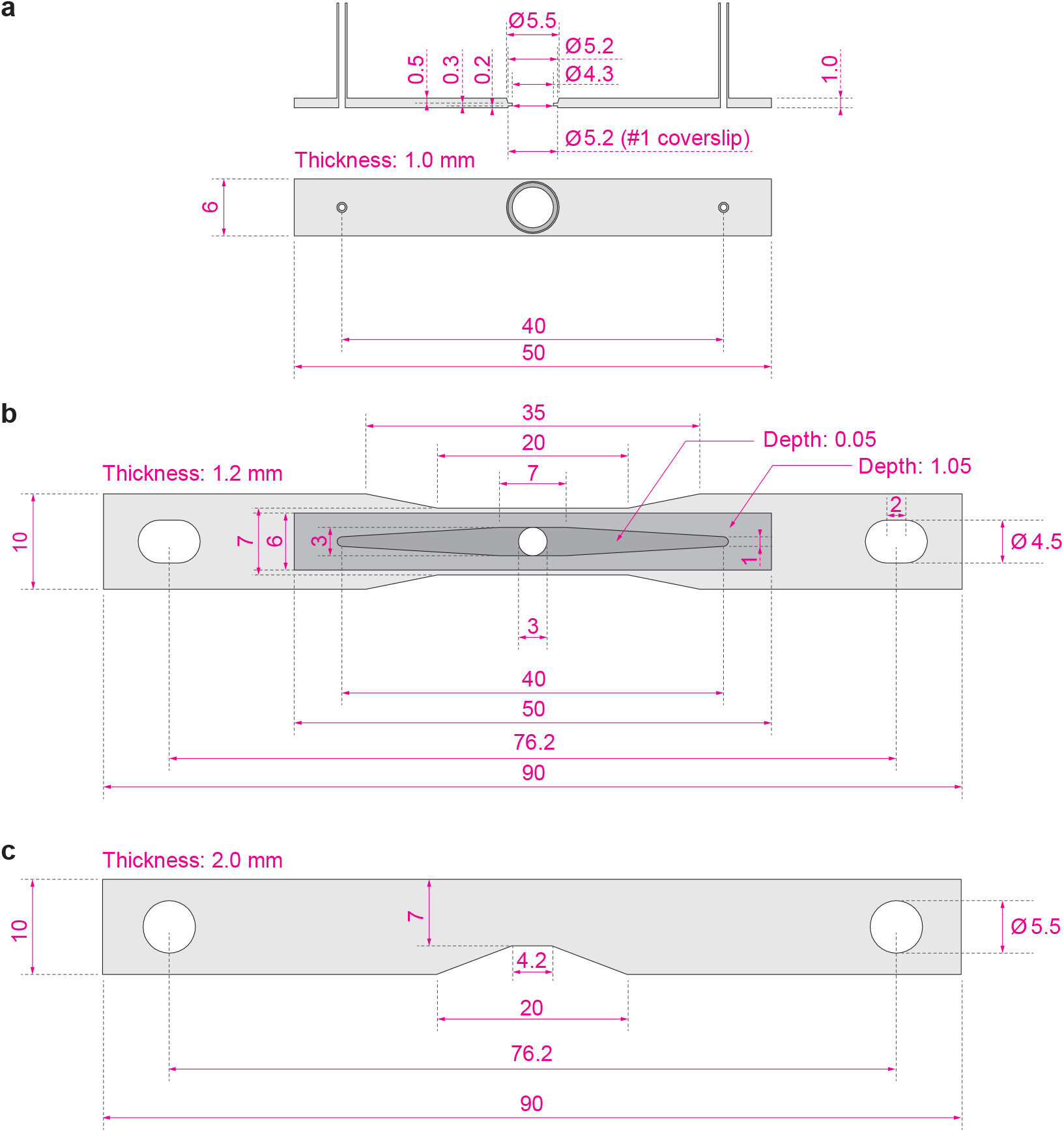
A detail drawing of a microfluidics-on-a-tongue device. (**a**) The upper piece for the top unit. It contains two connectors, one for multi-channel input and the other for output drainage. The open window is designed for mounting a coverslip with 5 mm in diameter and 0.15 mm in thickness. (**b**) The lower piece for the top unit, containing a fluidic channel and two thru-holes. The circular opening at the center of the channel is sealed when interfaced with the mouse tongue. (**c**) The bottom unit for gluing the tongue. All numbers without units are in mm.

**Supplementary Fig. 2.**
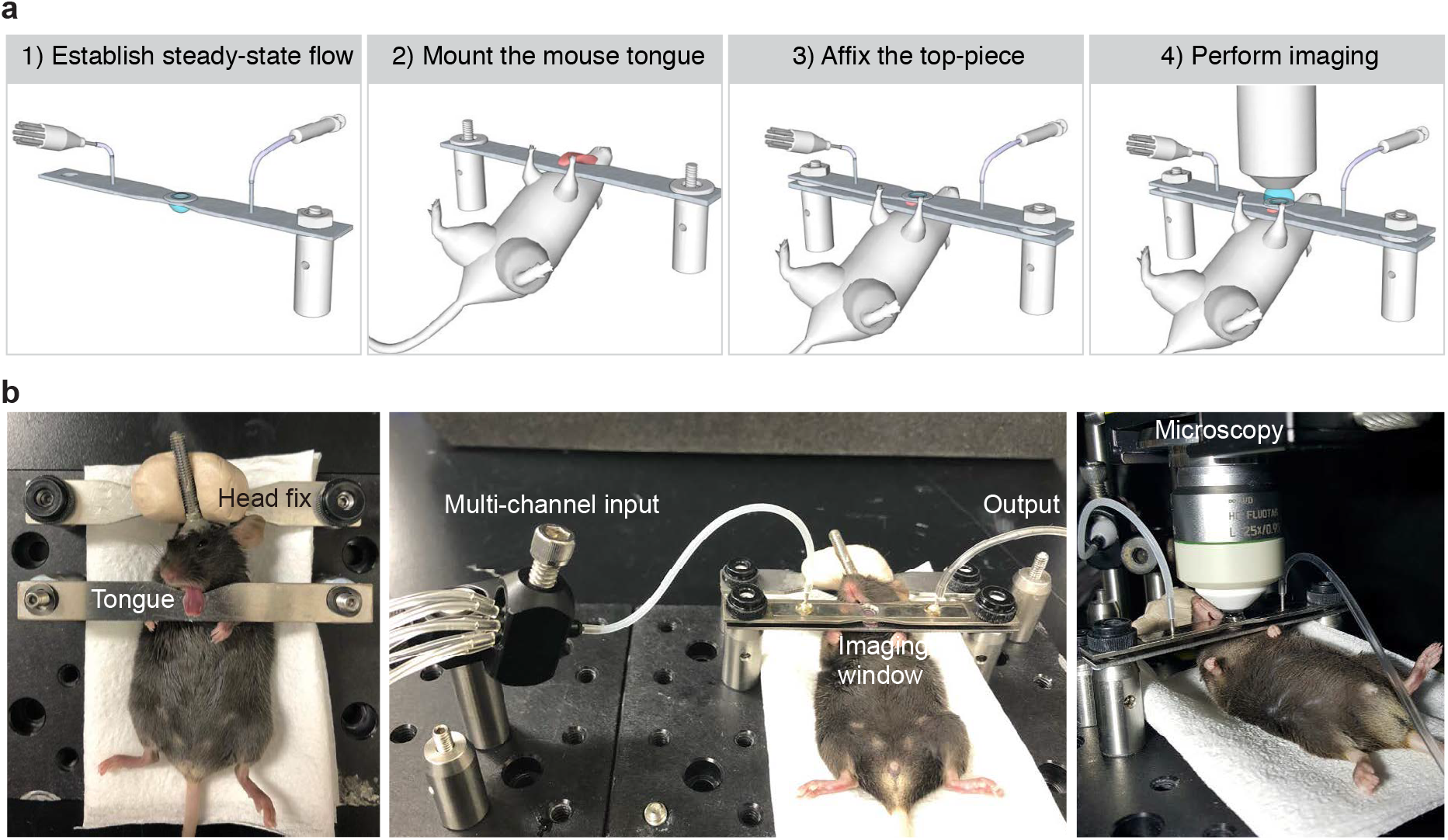
A step-by-step protocol for the preparation of functional taste cell imaging *in vivo*. (**a**) Mounting procedure. A microfluidic chamber is connected to the multi-channel input and the output to obtain quasi-steady-state flow (step 1). The ventral part of the tongue is glued onto the bottom plate (step 2). The top piece is placed on the dorsum of the tongue. The height is adjusted to ensure physiologic blood flow (step 3). Functional *in vivo* imaging on taste cells is performed under two-photon microscope (step 4). Detailed procedures are described in Online Methods. (**b**) Photographs of the preparations in (a). Left, the mouse tongue is attached to the bottom plate (left). Fluidic control system is assembled and mounted on the mouse tongue (middle). The preparation is mounted on an imaging system (right).

**Supplementary Fig. 3.**
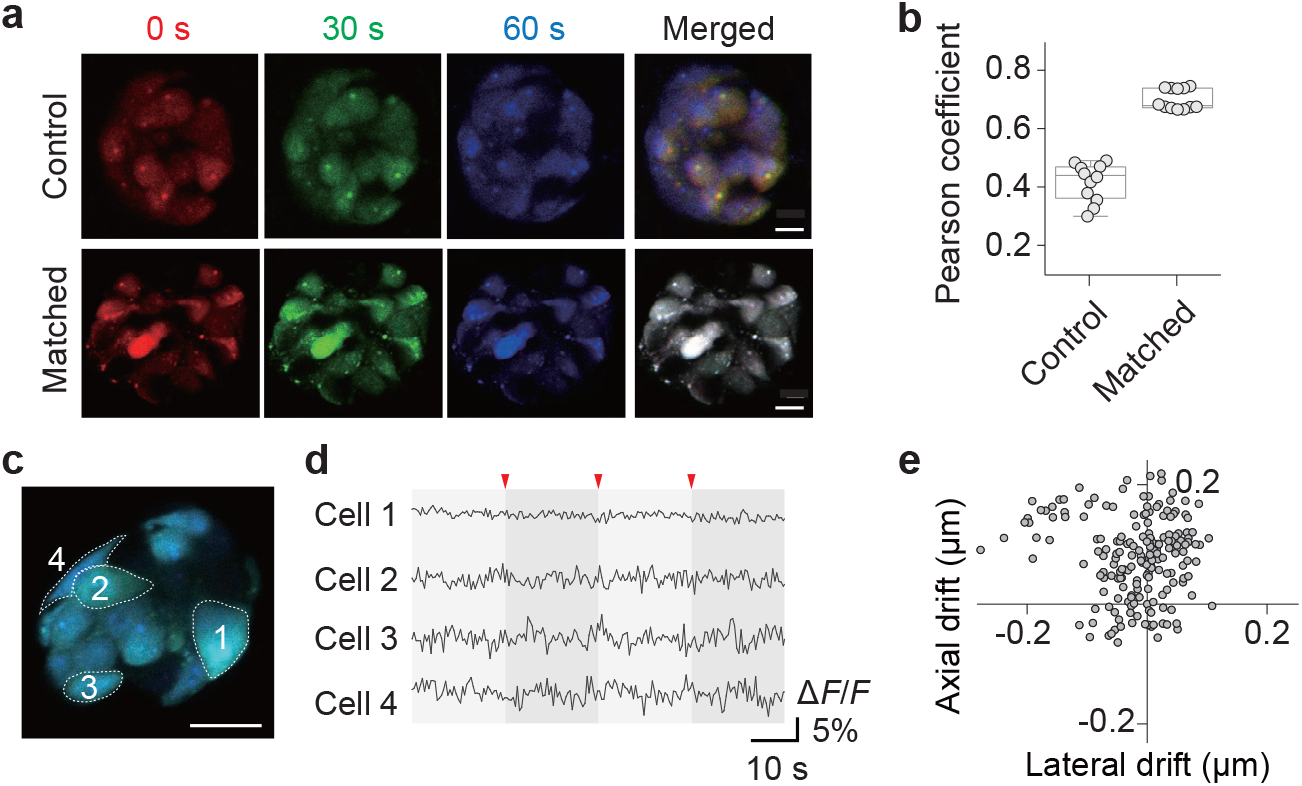
Imaging stability. (**a**) Time-series fluorescent imaging of a fungiform taste bud. The top row was acquired without matching input and output fluidics (control) and the bottom row was acquired after attaining quasi-steady-state flow (stabilized). Fluorescent intensity is pseudo-colored to highlight motion artifact in the merged image (the white color in the merged image represents no motion). Scalebar, 5 μm. (**b**) Motion artifact quantified by Pearson’s correlation coefficient of time-series images in (a). (**c,d**) Representative cellular calcium traces with switching fluidic channels (indicated by arrows). Scalebar, 10 μm. (**e**) Quantification of lateral and axial drifts of fungiform taste bud over 80 s. Each point indicates distance calculated from the first frame marked as an origin of the coordinates^2^.

**Supplementary Fig. 4.**
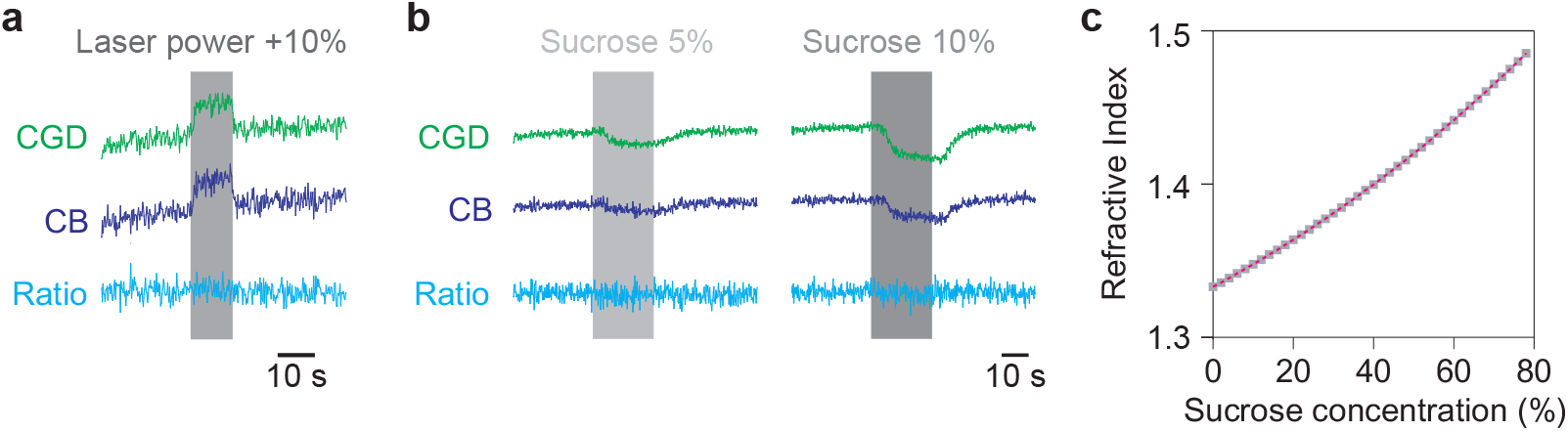
Verification of ratiometric analysis. (**a**) Ratiometric correction of laser fluctuation. Laser power was increased by 10% for duration of 10 s (indicated by gray). Note that ratiometric analysis effectively removed the artifact. (**b**) Ratiometric correction of tastant-induced optical artifact. Intensity fluctuation by introducing sucrose solution (5% and 10%) in a fluidic system was effectively corrected by the ratiometric analysis. (**c**) Refractive indices of sucrose solution in different concentration.

**Supplementary Fig. 5.**
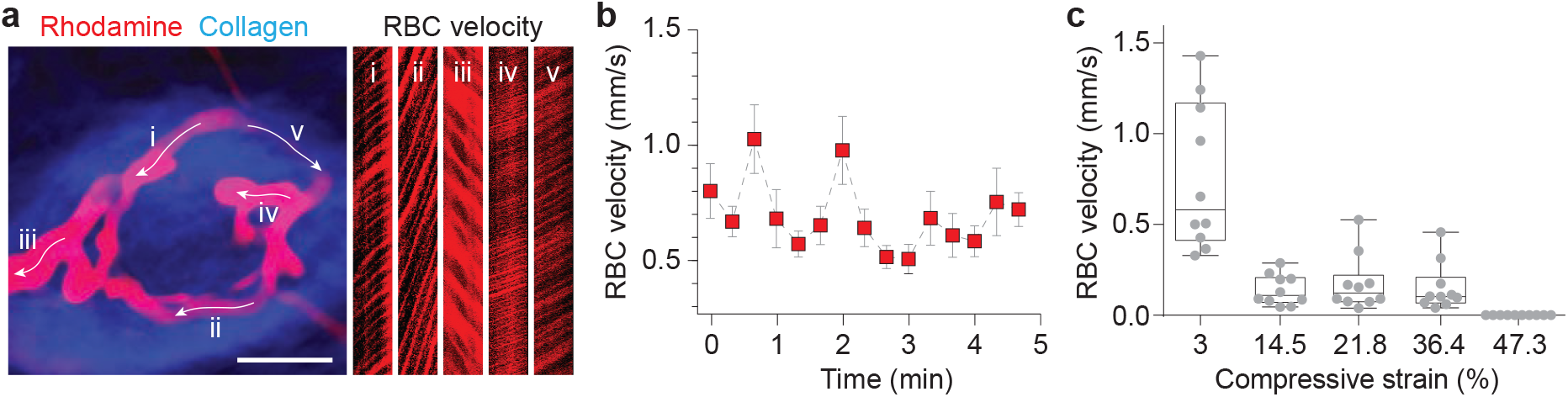
Characteristic of blood perfusion. (**a**) Quantification of blood flow. Rhodamine-B dextran (70 kDa) was intravenously injected and a fungiform taste bud was imaged by two-photon microscopy. The arrows indicate direction of the blood flow. RBC velocity was measured by the line scanning for each vascular segment (x = 20 μm, t = 150 ms). i: 0.43±0.09 mm/s, ii: 0.25±0.05 mm/s, iii: 0.727±0.13 mm/s, iv: 1.28±0.19 mm/s, v: 1.13±0.21 mm/s (n = 10 measurements). Scalebar, 30 μm. (**b**) Time-series measurement of blood flow. (**c**) Blood flow with respect to compressive strain. Strain is calculated based on the measured thickness of the mouse tongue (1.65±0.06 mm).

**Supplementary Fig. 6.**
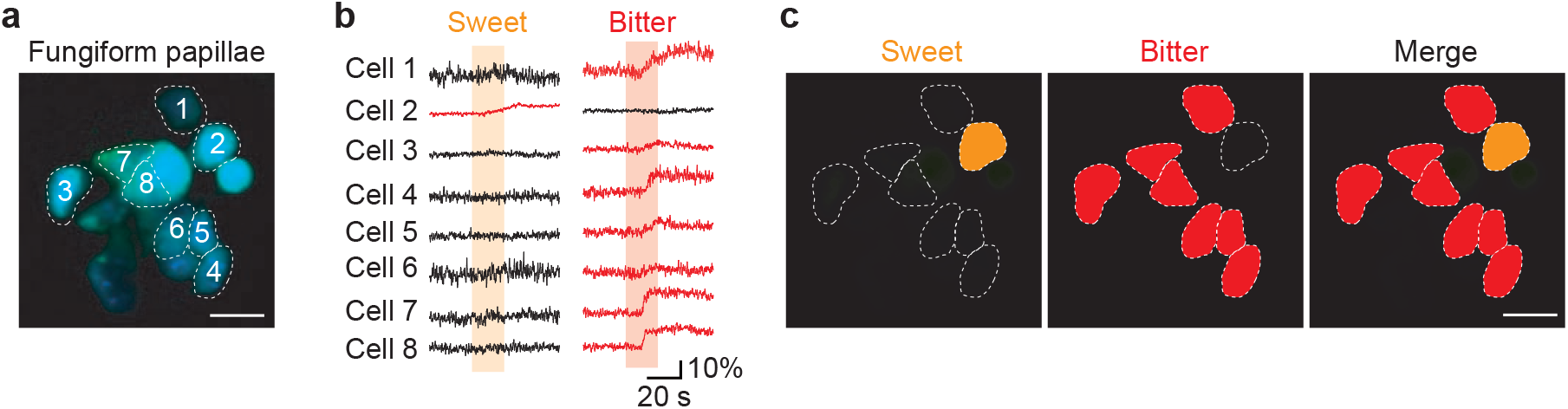
A bitter-tuned taste bud. (**a,b**) Functional activity of a bitter-tuned taste bud. Calcium responses to sweet and bitter tastants are shown (no activity observed in salty, sour, and umami). Note that 7 out of 8 taste cells responded to bitter. (**c**) A taste map for (a). Scalebar, 10 μm.

